# Membrane curvature sensing by model biomolecular condensates

**DOI:** 10.1101/2023.04.05.535714

**Authors:** Midhun Mohan Anila, Rikhia Ghosh, Bartosz Różycki

## Abstract

Biomolecular condensates (BCs) are fluid droplets that form in biological cells by liquid-liquid phase separation. Their major components are intrinsically disordered proteins. Vast attention has been given in recent years to BCs inside the cytosol and nucleus. BCs at the cell membrane have not been studied to the same extent so far. However, recent studies provide increasingly more examples of interfaces between BCs and membranes which function as platforms for diverse biomolecular processes. Galectin-3, for example, is known to mediate clathrin-independent endocytosis and has been recently shown to undergo liquid-liquid phase separation, but the function of BCs of galectin-3 in endocytic pit formation is unknown. Here, we use dissipative particle dynamics simulations to study a generic coarse-grained model for BCs interacting with lipid membranes. In analogy to galectin-3, we consider polymers comprising two segments – one of them mediates multivalent attractive interactions between the polymers, and the other one has affinity for association with specific lipid head groups. When these polymers are brought into contact with a multi-component membrane, they spontaneously assemble into droplets and, simultaneously, induce lateral separation of lipids within the membrane. Interestingly, we find that if the membrane is bent, the polymer droplets localize at membrane regions curved inward. Although the polymers have no particular shape or intrinsic curvature, they appear to sense membrane curvature when clustered at the membrane. Our results indicate toward a generic mechanism of membrane curvature sensing by BCs involved in such processes as endocytosis.

## 1 Introduction

Liquid-liquid phase separation (LLPS) underlies the formation of membraneless organelles such as stress granules and nucleoli [2]. The list of cellular compartments reported to be formed *via* LLPS is growing rapidly and touches a myriad of cell functions. In addition to membraneless organelles, other types of biomolecular condensates (BCs) are formed in the process of LLPS; examples include heterochromatin [26, 53], the transport channel in the nuclear pore complex [46], and membrane receptor clusters at the cell membrane [54].

The main macromolecular components of BCs are intrinsically disordered proteins (IDPs), which have no tertiary structure and exhibit large conformational fluctuations in aqueous solution [38, 41]. Instead of folding into one specific native structure, any IDP exhibits diverse conformations at native conditions [7, 20]. Since many of these conformations can be spatially extended, IDPs are able to bind multiple macromolecules *via* multivalent binding sites, even at low concentrations, which is a driving force for the formation of BCs [56].

Spectacular advances have been recently made in the research on BCs at the cell membrane. For example, the discovery of LLPS of zonula occludens proteins have explained how tight junctions are formed and how they function in mechanochemical signaling [6, 47]. LLPS has been shown to promote assembly of transmembrane proteins together with their cytoplasmic binding partners into (sub)micrometer-size clusters [54, 3, 58]. Consequently, LLPS has been implicated in the formation and functioning of clusters of various transmembrane receptors, including immune receptors, cell adhesion receptors, Wnt receptors, and glycosylated receptors [8]. BCs at the cell membrane have been recently shown to function in actin assembly [9] and in a Ras signaling pathway [22]. Evidence of membrane remodeling by BCs has been provided for the N-terminal low-complexity domain of fused in sarcoma (FUS LC) [57], for endocytic coat proteins with intrinsically disordered prion-like domains [5, 12], for transmembrane adaptor Linker of Activation of T-cells (LAT) during T-cell activation [55], for lipid vesicles within a synapsin-rich liquid phase [36], and for TIS granules interacting with the endoplasmic reticulum [33]. Such membrane reorganization processes mediated by BCs are anticipated to be a consequence of capillary forces generated at the membrane-condensate interface [16, 45]. Taken together, interfaces between BCs and membranes seem to provide platforms for diverse biomolecular processes, which are only beginning to be studied and understood.

An interesting example of a protein forming BCs and interacting with cell membranes is galectin-3. It is involved in numerous intra- and extra-cellular processes [39]. In particular, galectin-3 mediates glycosphingolipid-dependent biogenesis of clathrin-independent carriers [25]. Although oligomerization of galectin-3 has been implicated in this process, the molecular mechanisms underlying the endocytic pit formation remain elusive. However, the recent discovery that galectin-3 can undergo LLPS sheds new light on how this protein can perform its biological functions [10].

Galectin-3 is relatively small, soluble protein that comprises a carbohydrate recognition domain (CRD) and an intrinsically disordered N-terminal domain (NTD). This molecular architecture enables galectin-3 to bind many ligands and to form oligomers [28, 24, 31]. The CRD binds specifically *β*-galactosides, which mediates the binding of galectin-3 to various glycolipids in the cell membrane [31, 11]. The NTD mediates oligomerization of galectin-3 and, importantly, multivalent interactions between the NTDs drive the LLPS of galectin-3 [10].

Dissipative particle dynamics (DPD) simulations have been widely used to study various biomembrane processes, including membrane fusion [50], membrane rupture and damage [18], formation of lipid membrane domains [4], spontaneous curvature generation mechanisms [42, 43, 15], and membrane engulfment of liquid nano-droplets [44]. Here we employ DPD simulations to study a generic coarse-grained model for BCs interacting with lipid membranes. Specifically, by analogy to galectin-3, we consider polymers comprising two segments: The first segment mediates multivalent attractive interactions between the polymers, whereas the second one has affinity for glycolipid head groups. When these polymers are brought in contact with a two-component lipid membrane, they coalesce to form a nano-droplet and, simultaneously, associate with glycolipid head groups. These processes together induce lateral separation of lipids in the membrane. Interestingly, we discover that if the membrane is bent, or wavy, the polymer nano-droplet localizes on top of membrane regions that are concave. Although the polymers have no particular shape or intrinsic curvature, they appear to sense membrane concavity when clustered at the membrane. Since the coarse-grained model that we employ in this study is generic and not uniquely parameterized for specific IDPs and lipids, we argue that the membrane curvature sensing may be a general, geometric feature of BCs involved in such processes as endocytosis.

The sensing of membrane curvature by the polymer nano-droplets resembles the capillary effect of curvotaxis, i.e. spontaneous motion of fluid droplets towards more concave, or less convex, regions of a substrate [32, 14, 34]. The curvotaxis of a fluid droplet on a solid substrate can be explained simply by considering how interfacial energies depend on the local shape of the substrate surface. Therefore, to better understand the DPD simulation results on membrane curvature sensing, we put forward a toy model based on the theory of wetting. The toy model not only captures the effect of curvature sensing but also helps to understand the underlying physics. Moreover, it allows us to tell apart the effects of curvature sensing and curvature generation.

This article is organized as follows. In section 2 we give a detailed description of the coarse-grained model for the polymers and lipids, introduce the equations of motion that govern the coarse-grained dynamics, and describe simulation data analysis methods. In section 3 we describe our results of coarse-grained simulations of three systems: (i) the two-component membrane in the absence the polymers (section 3.1), (ii) the polymers in solution, i.e. in the absence of the membrane (section 3.2), and (iii) the two-component membrane interacting with the polymers (sections 3.3 and 3.4). Then we present the toy model that captures the sensing of membrane curvature by liquid droplets (section 3.5). In section 4 we explain the mechanism of membrane curvature sensing that emerges from our simulations and calculations, and discuss implications of our results.

## 2 Methods

We use an explicit-solvent coarse-grained model with dissipative particle dynamics [19, 49] to simulate systems composed of several thousand lipids and a few hundred IDPs on time scales up to 200 *µ*s. The model membrane consists of two types of lipids, which we denote here by lipid A and lipid B. Lipid A has two hydrocarbon tails, each represented by a chain of 6 hydrophobic beads *T*, and a head group composed of 3 hydrophilic beads *H*_*A*_. Lipid B is composed of the same hydrophobic tails as lipid A and has a larger head group, which is composed of 4 hydrophilic beads *H*_*B*_, to model the bulky head group of glycosphingolipids. The lower monolayer of the membrane contains only lipids of type A. The upper monolayer of the membrane, on the other hand, contains 60% of lipid A and 40% of lipid B. Each of the monolayers comprises the same total number of lipids. A molecule of galactin-3 is modeled here by a linear polymer P composed of 32 beads of type *P*_*T*_, representing the intrinsically disordered NTD, followed by 4 beads of type *P*_*C*_, representing the CRD. Water is represented by single beads of type *W*. The different types of beads and molecules incorporated in the coarse-grained model are summarized in figure 1 (where water beads *W* are shown in pale blue; lipid head-group beads *H*_*A*_ and *H*_*B*_ in tan and yellow, respectively; lipid tail beads *T* in gray; and polymer cap and tail beads, *P*_*C*_ and *P*_*T*_, in cyan and blue, respectively).

**Figure 1:**
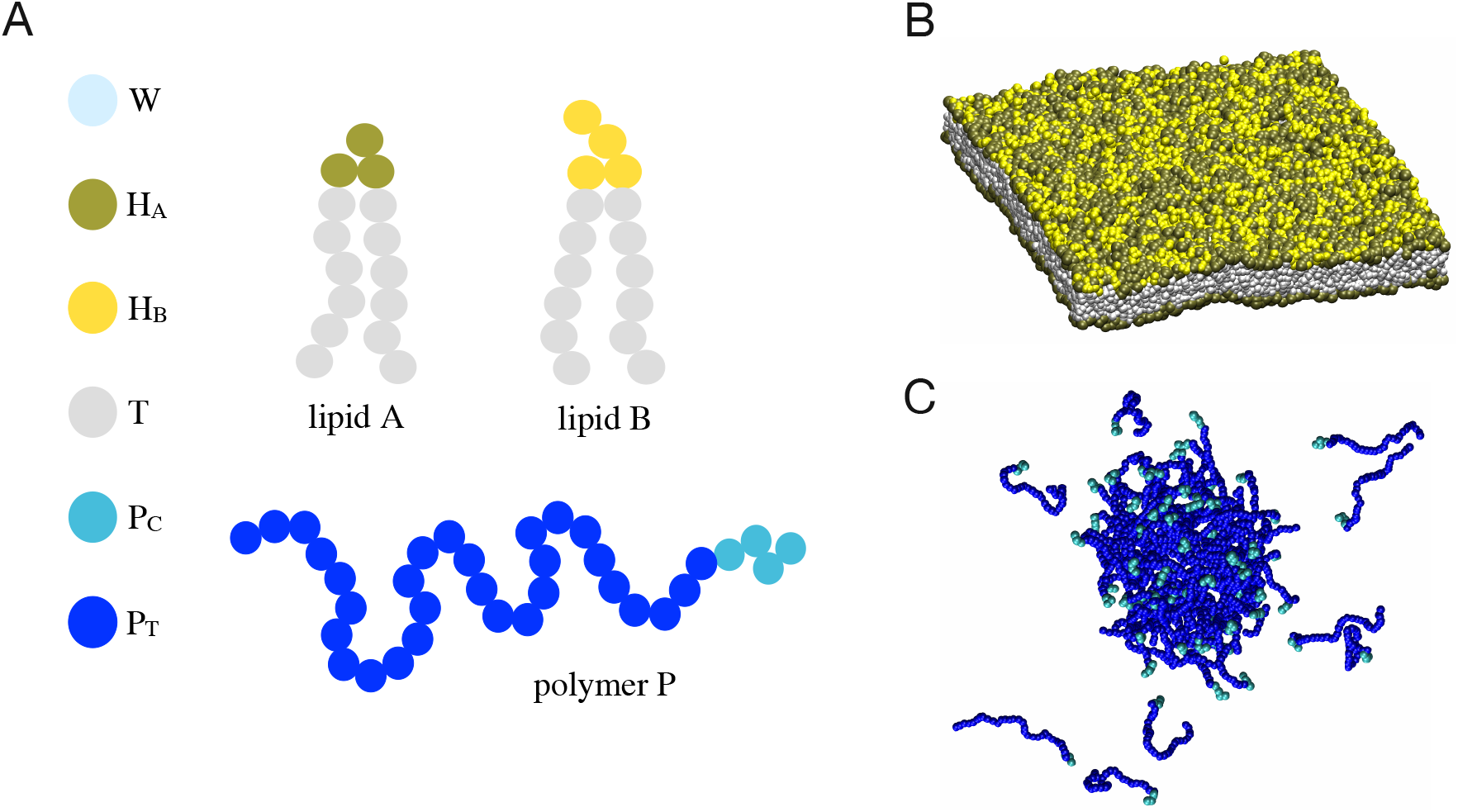
(A) Schematic diagram of the different bead types and molecules used in the coarse-grained simulations. The head-group beads *H*_*A*_ (tan) and *H*_*B*_ (yellow) of lipid A and lipid B, respectively, have similar molecular features except for their tendency to associate differently to the polymer caps formed of *P*_*C*_ beads (cyan). The tail beads *T* (gray) of lipid A and lipid B represent the same molecular units. The polymer consists of a disordered tail, which is composed of 32 beads of type *P*_*T*_ (blue), and a cap composed of 4 beads of type *P*_*C*_ (cyan) that selectively associate with the head group of lipid B. (B) Simulation snapshot of a lipid membrane. Water beads *W* are not shown for clarity. The upper monolayer of the membrane is composed of lipid A (head groups in tan) and lipid B (head groups in yellow) in 4:1 ratio. The lower monolayer (not visible in this view) contains lipid A only. (C) Simulation snapshot of a spontaneously formed nano-droplet of polymers P. Water beads *W* are not shown for clarity. The self-assembly of the polymers is mediated by the tail beads *P*_*T*_.

In DPD each pair of beads interacts *via* three short-ranged, non-bonded, additive forces: (1) random force 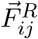 (2) dissipative force 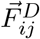 and (3) conservative force 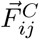. The random force

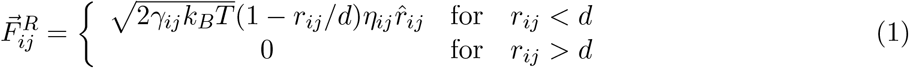

represents thermal noise. Here, *k*_*B*_ is the Boltzmann constant, *T* is the absolute temperature, *γ*_*ij*_ is the friction coefficient, and *η*_*ij*_ represents the Gaussian white noise with average value *η*_*ij*_(*t*) = 0 and auto-correlation function *η*_*ij*_(*t*)*η*_*i*_ *j* (*t*) = *δ*_*ii*_ *δ*_*jj*_ *δ*(*t* − *t*). The unit vector 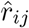 points from bead *j* to bead *i*, and *r*_*ij*_ denotes the distance between the two beads. The dissipative force

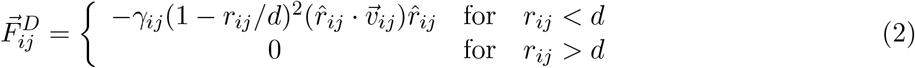

acting between beads *i* and *j* depends on the relative velocity 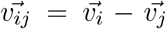 of the beads. The conservative force

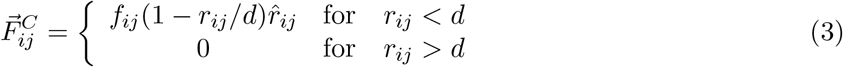

defines the identity of the beads (e.g. their level of hydrophobicity or hydrophilicity) because the force parameters *f*_*ij*_ *>* 0 determine the strength of repulsion between beads *i* and *j*. The numerical values of the force parameters *f*_*ij*_ that we use for each pair of beads in our model is given in Table 1.

**Table 1:** DPD force parameters *f*_*ij*_ in the units of *k*_*B*_*T/d*

| $f_{ij}$ | $j = W$ | $j = T$ | $j = H_A$ | $j = H_B$ | $j = P_T$ | $j = P_C$ |
| --- | --- | --- | --- | --- | --- | --- |
| $i = W$ | 25 | 75 | 30 | 30 | 25 | 25 |
| $i = T$ | 75 | 10 | 50 | 50 | 50 | 50 |
| $i = H_A$ | 30 | 50 | 30 | 30 | 30 | 30 |
| $i = H_B$ | 30 | 50 | 30 | 30 | 30 | 4 |
| $i = P_T$ | 25 | 50 | 30 | 30 | 18 | 25 |
| $i = P_C$ | 25 | 50 | 30 | 4 | 25 | 25 |

For lipid A we use the six-bead-per-tail model and the corresponding set of parameters *f*_*ij*_ introduced in our previous work [42, 43]. Lipid B and lipid A are characterized by almost identical sets of parameters *f*_*ij*_ except for *f*_*H*_*A,P*_*C*_ = *f*_*HB*_,*P*_*C*_, which distinguishes their head-group interactions with the polymer caps. The choice of *f*_*HB*_,*P*_*C*_ = 4*k*_*B*_*T/d* and *f*_*HA*_,*P*_*C*_ = 30*k*_*B*_*T/d* ensures that the polymer cap (representing the CRD of galection-3) preferentially associates with the head group of lipid B (representing glycosphingolipids) and not with the head group of lipid A, whereas the bulkier head of lipid B (composed of four beads of type *H*_*B*_) strengthens the effective attraction between the polymer cap and the lipid-B head. On the other hand, the lipid head-group interaction parameters *f*_*HA*_,*H*_*A*_ ≠ *f*_*HB*_,*H*_*B*_ = *f*_*HA*_,*H*_*B*_ = 30*k*_*B*_*T/d* are chosen to prevent spontaneous separation of lipids A and B in the membrane. The choice of *f*_*PT*_, *P*_*T*_ = 18*k*_*B*_*T/d* is made to bring about association of the polymer tails that represent the intrinsically disordered N-terminal domain of galectin-3. We note that IDPs have been modeled previously in DPD simulations as linear polymers with self-associating endcaps [51]. This model, however, is fundamentally different from our current model, where the self-association is mediated by the entire disordered tail.

In addition to the non-bonded forces 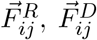 and 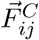, the adjacent beads within the lipids and polymers are bound together by a harmonic potential

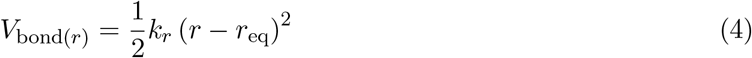

with a spring constant *k*_*r*_ = 128*k*_*B*_*T/d*^2^ and the equilibrium separation *r*_eq_ = *d/*2. Additionally, a bending potential

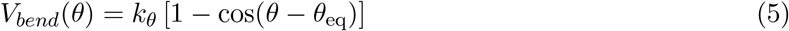

is implemented to stiffen the chains of the lipids and polymers. The bending potential acts between two consecutive bonds along a chain. For the bond pairs in the lipids (*T* − *T* − *T, T* − *T* − *H*_*A*_ in lipid A and *T* − *T* − *T, T* − *T* − *H*_*B*_ in lipid B), the bending constant *k*_*θ*_ = 15*k*_*B*_*T* and the equilibrium value of the bond angle *θ*_eq_ = 0 (meaning collienar bonds) [49]. In order to represent the disordered nature of the polymer tails, we choose the corresponding bending constant between tail bead pairs *P*_*T*_ − *P*_*T*_ − *P*_*T*_ relatively soft, *k*_*θ*_ = 5*k*_*B*_*T* [48].

All beads have the same diameter *d* and the same mass *m*. These two quantities provide the basic units of length and mass, respectively. The basic unit of energy is chosen to be the thermal energy, *k*_*B*_*T*. Therefore, the basic time scale is *τ* = (*d*^2^*m/k*_*B*_*T*)^1*/*2^. It has been estimated in earlier studies that *d ≈* 0.8 nm and *τ ≈* 1 ns [17, 42, 43].

We performed the DPD simulations in the (*N, V, T*) ensemble using the Osprey-DPD package with the simulation time step ∆*t* = 0.02*τ* [17], the bead density of 3*/d*^3^ as typically in DPD simulations [19, 49], and a cubic box of size *L* = 48 *d ≈* 40 nm. We simulated systems with different values of the projected area per lipid

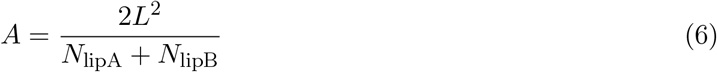

Here, *N*_lipA_ and *N*_lipB_ denote the number of lipids A and B, respectively, and the membrane spans the simulation box laterally in all of the simulation systems.

We constructed the membrane in such a way that the number of all lipids in the upper monolayer was 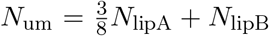, the number of lipids in the lower monolayer was 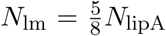 and the two numbers were equal, *N*_um_ = *N*_lm_, implying *N*_lipA_ = 4 *N*_lipB_. Thus the molar ratio of lipid A and lipid B in the upper monolayer was 3:2.

The DPD simulation trajectories were visualized using VMD [23]. In the analysis of the simulation trajectories we quantified the amount of contacts between beads of specific types using the coordination number

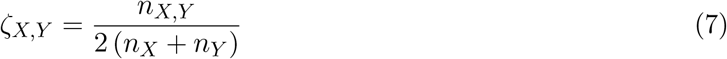

Here, *n*_*X,Y*_ denotes the number of inter-molecular contacts between all the beads of type X and of type Y, and *n*_*X*_ is the number of beads of type X in the system. We assume here that a pair of beads is in contact if the distance between their centers is smaller than or equal to the bead diameter *d*.

A decrease of *ζ*_*X,Y*_ with time means that more inter-molecular contacts between beads X and beads Y are broken than formed. On the other hand, an increase of *ζ*_*X,Y*_ with time implies a growth of inter-molecular contacts between beads of type X and of type Y, which may indicate formation of multimers or condensates of molecules containing these beads.

In order to identify polymer assemblages and quantify formation of BCs in the system under study, we applied agglomerative clustering to configurations obtained from the DPD simulations. At the beginning of the clustering procedure, each of the polymers is taken as a single cluster. Then clusters are merged successively. Two clusters get merged if any polymer from one cluster makes at least one inter-molecular contact with any polymer from the other cluster. The clustering procedure ends when no more clusters can be merged. Two useful quantities that result from the clustering procedure are 1) the number *n*_*C*_ of clusters comprising more than one polymer, and 2) the fraction *f*_LC_ of polymers in the largest cluster (LC), i.e. the number of polymers forming the LC divided by the total number of polymers in the system.

## 3. Results

### 3.1 The mixture of lipid A and lipid B in the membrane is homogeneous in the absence of interactions with the polymers

We performed DPD simulations of lipid bilayers with the lower leaflet containing only lipids of type A and the upper leaflet composed of lipid A and lipid B in 3:2 ratio. The polymers were not included in the simulation system. We performed the simulations for different values of the projected area per lipid, *A*, as defined by equation (6) and ranging from 1.04 *d*^2^ to 1.31 *d*^2^. For large values of *A*, the lipid bilayers are planar and stretched. Smaller values of *A* correspond to rough or undulating bilayers.

Figure 2 shows the coordination numbers *ζ*_*HA*_,*H*_*A*_, *ζ*_*HB*_,*H*_*B*_ and *ζ*_*HA*_,*H*_*B*_ for the lipid head-group beads, *H*_*A*_ and *H*_*B*_, as a function of time. The coordination numbers are defined by equation (7) and quantify the amount of contacts between beads of different types in the simulation system. The different colors in figure 2 indicate different values of the projected area per lipid (data for *A* = 1.04 *d*^2^ are shown in blue; for *A* = 1.09 *d*^2^ in red; for *A* = 1.23 *d*^2^ in black; and for *A* = 1.31 *d*^2^ in brown). The coordination numbers *ζ*_*HA*_,*H*_*A*_, *ζ*_*HB*_,*H*_*B*_ and *ζ*_*HA*_,*H*_*B*_ remain practically constant in time and independent of the projected area per lipid. We also checked that the lipids did not flip between the upper and lower leaflets during the simulation runs [42]. These results together show that the mixture of lipid A and lipid B in the upper leaflet remains homogeneous throughout the simulation time of 30 *µ*s.

**Figure 2:**
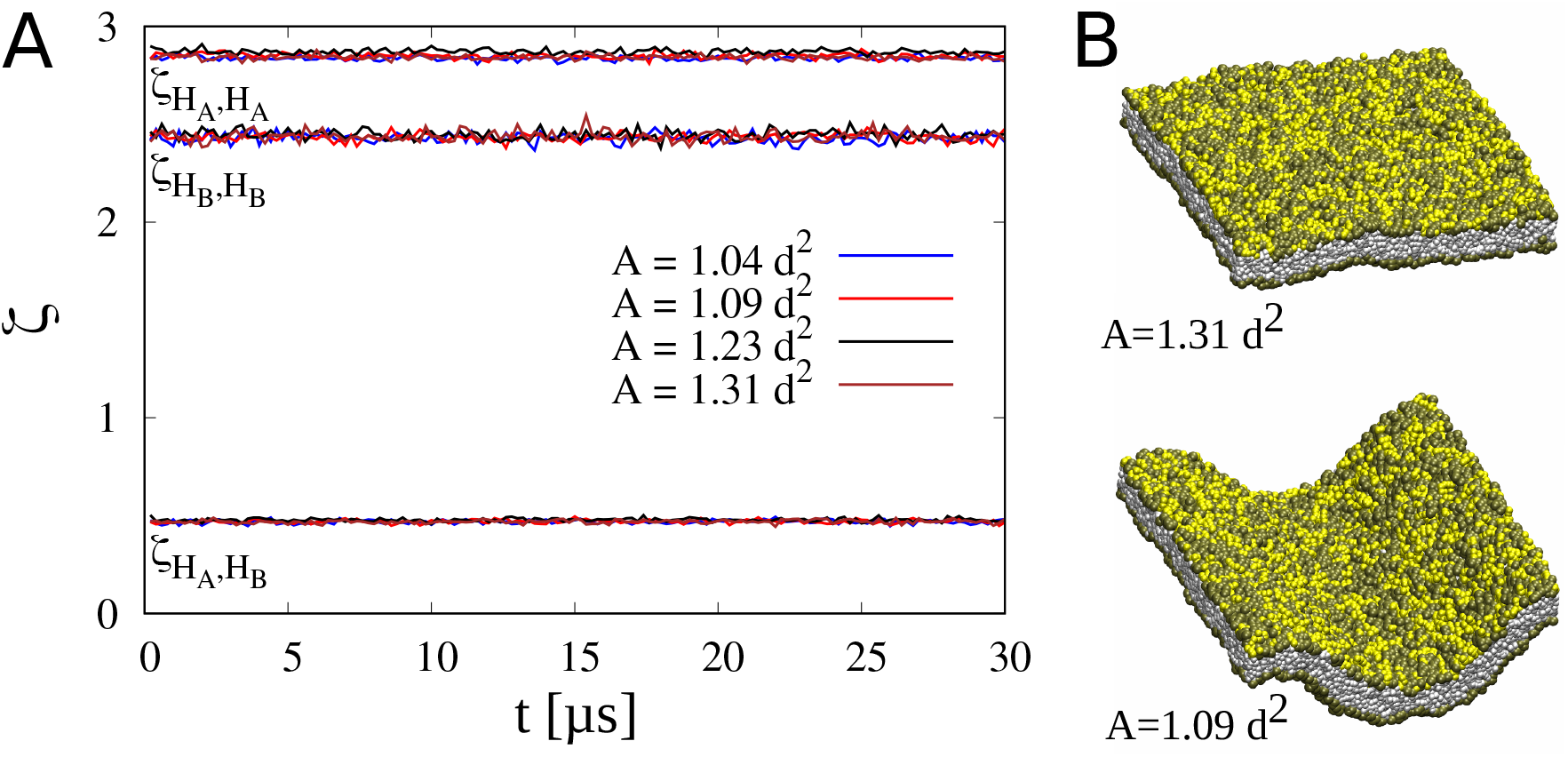
Lipids in the upper leaflet form a homogeneous mixture in the absence of interactions with the polymers. (A) Coordination numbers *ζ*_*HA*_,*H*_*A*_, *ζ*_*HB*_,*H*_*B*_ and *ζ*_*HA*_,*H*_*B*_ for the lipid headgroup beads, *H*_*A*_ and *H*_*B*_, as a function of time obtained from the DPD simulations of lipid bilayers in the absence of interactions with the polymers. The different colors indicate different values of the projected area per lipid (data for *A* = 1.04 *d*^2^ are shown in blue; for *A* = 1.09 *d*^2^ in red; for *A* = 1.23 *d*^2^ in black; and for *A* = 1.31 *d*^2^ in brown). (B) Two snapshots of lipid bilayers obtained from the DPD simulations with *A* = 1.31*d*^2^ and *A* = 1.09*d*^2^. The color code is as in figure 1, i.e. the lipid tails are shown in gray and the head groups of lipid A and lipid B are in tan and yellow, respectively. The water beads *W* are not shown.

### 3.2 At sufficient concentrations, the polymers in solution undergo LLPS in the absence of interactions with the lipids

We next performed DPD simulations of polymer P in solution with the molar fraction *φ* of the polymer ranging from 2 *·* 10^−4^ to 2 *·* 10^−3^. No lipids were included in the simulation system. The spatial distribution of the polymers was uniform and random at the beginning of the simulation runs.

Figure 3A shows the coordination number *ζ*_*PT*_, *P*_*T*_ as a function of time for different molar fractions of polymer P (data for *φ* = 2 *·* 10^−4^ are shown in black; for *φ* = 4 *·* 10^−4^ in blue; for φ= 6 *·* 10^−4^ in brown; for *φ* = 1.2 *·* 10^−3^ in green; and for *φ* = 2 *·* 10^−3^ in red). The increase of *ζ*_*P T*_, *P*_*T*_ observed for *φ >* 2 *·* 10^−4^ means that, on average, more inter-molecular contacts between the polymer tails is formed than broken during the simulation time, which suggests progressive association of the polymers. Visual inspection of the system configurations reveals that the polymers spontaneously assemble into clusters, or nano-droplets, as depicted in 3B.

**Figure 3:**
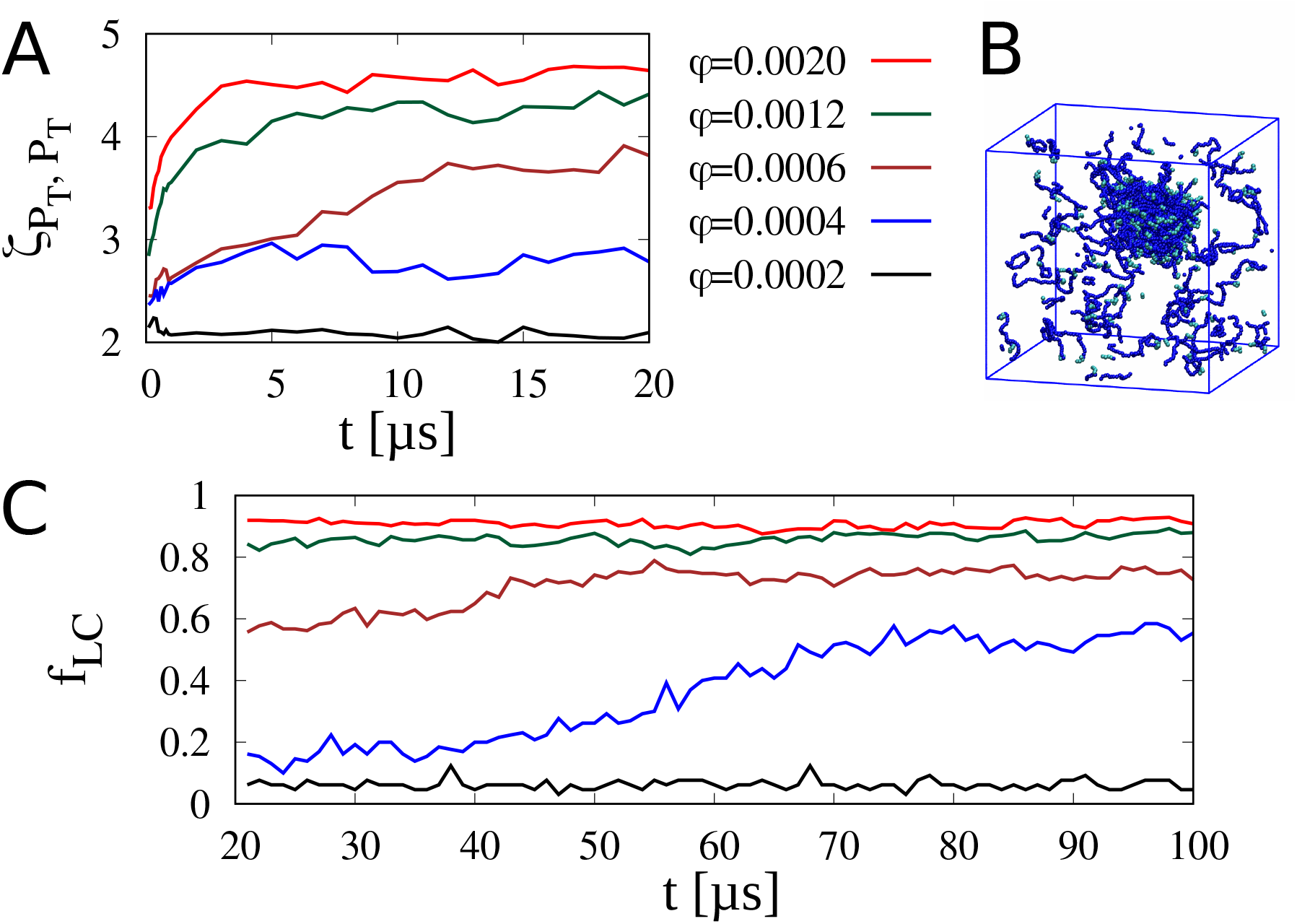
At sufficient molar fractions *φ*, the polymers in solution assemble spontaneously into fluid clusters or nano-droplets. (A) Time evolution of the coordination number *ζ*_*PT*_, *P*_*T*_ related to inter-molecular contacts between the polymer tails. Data obtained from the DPD simulations of polymer P in the absence of interactions with lipid A or lipid B. Different colors correspond to different molar fractions of the polymer (data for *φ* = 2 10^−4^ are shown in black; for *φ* = 4 10^−4^ in blue; for *φ* = 6 10^−4^ in brown; for *φ* = 1.2 10^−3^ in green; and for *φ* = 2 10^−3^ in red). (B) Simulation snapshot of a polymer cluster, or a nano-droplet, in equilibrium with single polymer chains. The color code is as in figure 1, i.e., the polymer tails are in blue and the polymer caps in cyan. The water beads *W* are not shown. (C) Fraction *f*_LC_ of polymers in the largest cluster as a function of time (color code as in panel A).

We extended the DPD simulations to 100 *µ*s and – in order to identify groups of polymers making mutual contacts – applied a clustering algorithm to the simulation system configurations. The clustering algorithm is described in the Methods section. Figure 3C shows the fraction *f*_LC_ of polymers forming the largest cluster (LC) as a function of time for different molar fractions *φ* (color code as in figure 3A). At *φ* = 2 *·* 10^−3^, i.e. at the largest molar fraction investigated in our simulations, *f*_LC_ is of the order of 0.9 during the simulation time, implying that a great majority of the polymers are assembled into one large cluster. Fluctuations of *f*_LC_ indicate that polymers associate with and dissociate from the LC during the simulation time, which suggests that the LC is in equilibrium with much smaller clusters or single polymers. At *φ* = 2 *·* 10^−4^, on the other hand, *f*_LC_ is about 0.1 or smaller during the simulation time, indicating that the system of polymers is in a dilute phase. At a somewhat larger molar fraction of *φ* = 4 *·* 10^−4^, however, *f*_LC_ increases slowly in time and reaches the value *f*_LC_ *≈* 0.6 at *t*=100 *µ*s, indicating a gradual condensation process. Taken together, we conclude that if *φ ≥* 4 *·* 10^−4^, the polymers condensate and form nano-droplets in solution.

Since the bead diameter *d ≈* 0.8 nm in our coarse-grained model [42] and, thus, the simulation box length *L* is about 40 nm, the molar fraction *φ* = 4 *·* 10^−4^ (at which the nano-droplet formation is observed in the DPD simulations) corresponds to the polymer concentration of about 3 mM. For comparison, at temperature *T* = 30 ^*?*^C and with no salt in solution, the NTD of galectin-3 undergoes LLPS at the concentration of 2.3 mM [10]. This degree of agreement between our simulation results and experimental observations gives grounds for our choice of the tail-tail interaction parameter *f*_*PT*_, *P*_*T*_ = 18*k*_*B*_*T/d* (Table 1).

We also computed the radius of gyration of single polymers, *R*_g_, at the smallest molar fraction *φ* = 2 *·* 10^−4^ at which the system is in the dilute state. We obtained 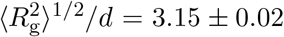 With *d ≈* 0.8 nm we estimate that 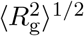 is of the order of 2.5 nm in our simulations, which compares quite well with the radius of gyration of galectin-3 of 2.9 nm as determined in small angle X-ray scattering experiments [31]. This comparison justifies the choice of the length and stiffness of polymer P in our coarse-grained model.

To check whether the polymer clusters are in a fluid state, we devised a simulation trajectory analysis method that resembles FRAP experiments. Namely, at time instance *t* = *t*_0_ we select all of the polymer beads with the Cartesian coordinates *x* and *y* that fulfill the condition |*x* − *L/*2| *< r*_c_ and |*y* − *L/*2| *< r*_c_ with *r*_c_ = 3 *d*. These beads are located inside a cuboid parallel to the *z*-axis and centered in the middle of the simulation box. We label these beads in red and all other polymer beads in green (figure 4, left hand-side panel). Then we take simulation snapshots at *t > t*_0_ and watch the location of the red beads (figure 4, central and right hand-side panels). Mixing the red and green beads demonstrates diffusion.

**Figure 4:**
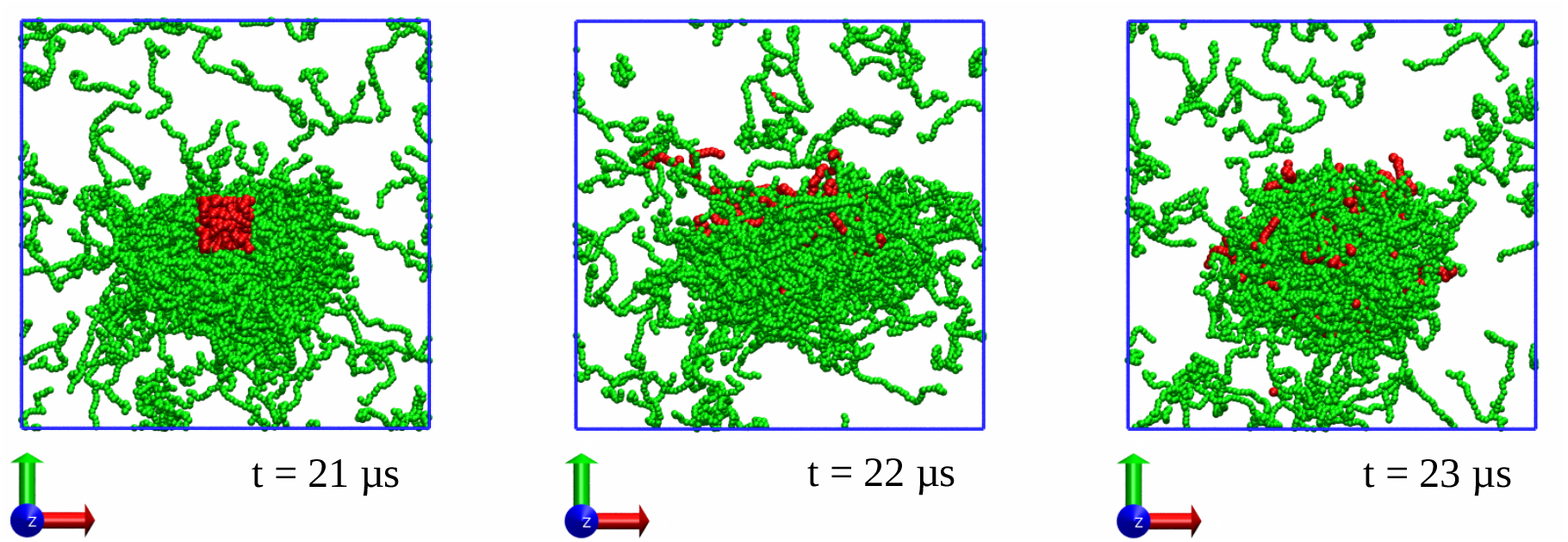
Simulation snapshots showing that the polymer system with φ = 1.2 10^−3^ is fluid. At the time instance *t*=21 *µ*s (snapshot on the left hand side), when the LC is already established, a group of beads is selected and marked in red. The red beads are located inside a cuboid parallel to the *z*-axis and centered in the middle of the simulation box. All other breads are marked in green. At *t*=22 *µ*s (snapshot in the middle) and *t*=23 *µ*s (snapshot on the right hand side), the red beads are mixed with the green beads, which demonstrates polymer diffusion and, thus, fluidity of the LC.

Figure 4 shows results of this analysis applied to a trajectory of the polymer system with φ = 1.2 *·* 10^−3^. Here, *t*_0_=21 *µ*s, when the LC is already formed. The snapshots at *t*=22 *µ*s and *t*=23 *µ*s show that the red beads diffuse throughout the LC, demonstrating that the polymer cluster is fluid.

### 3.3 Interactions between the cap of polymer P and the head group of lipid B induce lateral separation of lipids in the membrane

We next performed DPD simulations of the two-component membrane (with several different values of the area per lipid *A*) in the presence of polymers P (with molar fractions φ = 5 *·* 10^−4^ and φ = 10^−3^). The simulation systems were set up in such a way that initially, at time *t* = 0, the polymers were positioned randomly in the fluid above and below the membrane whereas the two types of lipids were distributed uniformly in the upper leaflet. In the course of the simulations we observed the polymers associating *via* their end caps with the lipid-B head groups and, simultaneously, assembling into clusters, which together led to lateral segregation of the two types of lipids within the upper leaflet of the membrane, see the simulation snapshots in figure 5E.

**Figure 5:**
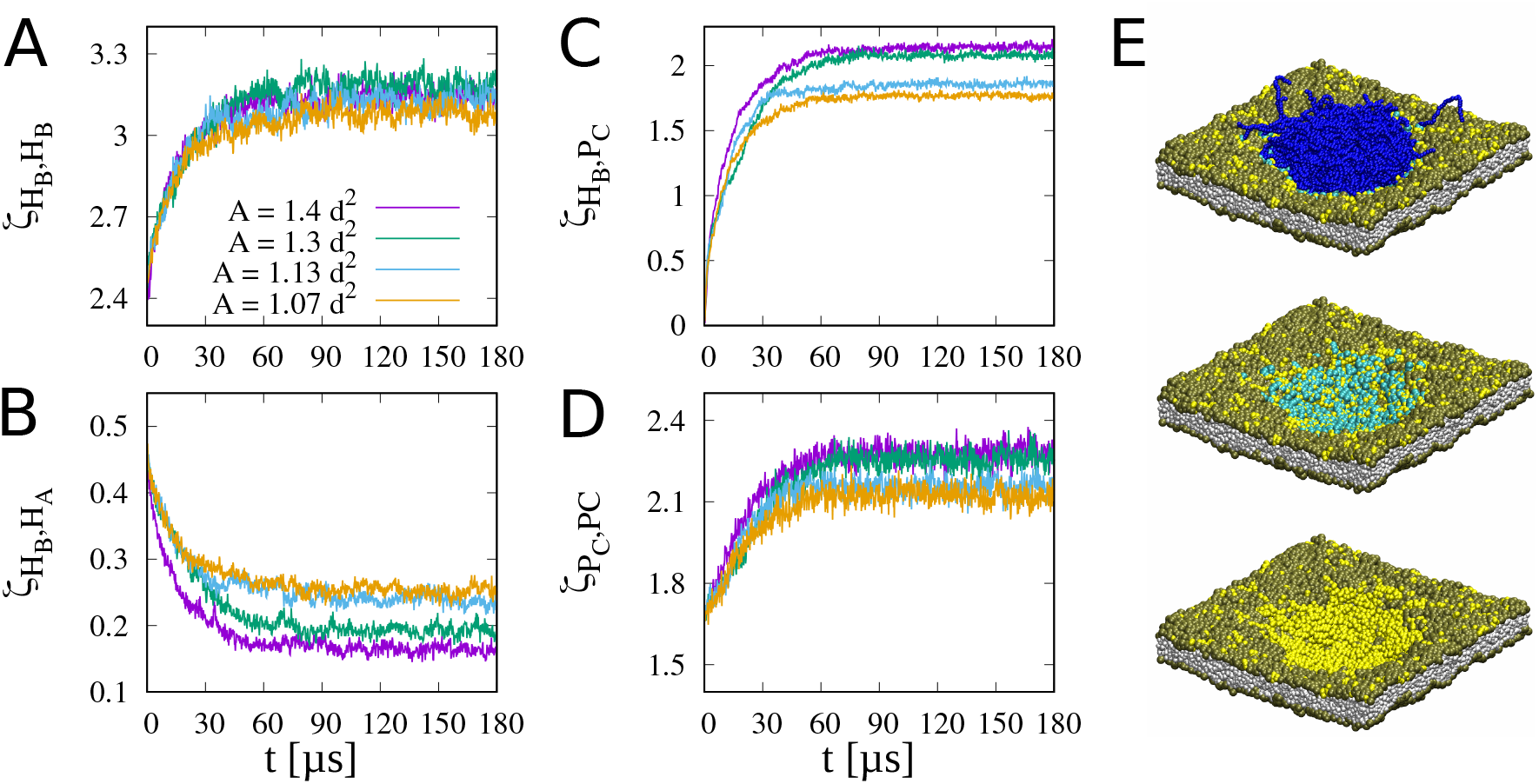
Interactions between the cap of polymer P and the head group of lipid B induce separation of lipid A from lipid B in the upper leaflet of the membrane. (A–D) Coordination numbers *ζ*_*HB*_,*H*_*B*_ (A), *ζ*_*HB*_,*H*_*A*_ (B), *ζ*_*HB*_,*P*_*C*_ (C) and *ζ*_*PC*_,*P*_*C*_ (D) as a function of time. Results obtained from the DPD simulations of the polymers in contact with the two-component membrane. The molar fraction of polymer P is φ = 5 10^−4^. The different colors indicate different values of the projected area per lipid (data for *A* = 1.4 *d*^2^ are shown in magenta; for *A* = 1.3 *d*^2^ in green; for *A* = 1.13 *d*^2^ in blue; and for *A* = 1.07 *d*^2^ in orange). (E) Snapshots of the simulation system taken at *t* = 180*µ* s. Here, *A* = 1.22 *d*^2^ and *φ* = 10^−3^. The color code is as in figure 1. The water beads *W* are not shown for clarity. The upper snapshot shows that polymers P form a nano-droplet at the membrane. In the middle panel, the polymer tails are not displayed. It can be seen that the polymer caps (cyan) co-localize with the lipid-B head groups (yellow). In the bottom snapshot, only the lipids are shown. It can be seen that lipid A (tan) segregate from lipid B (yellow).

Figure 5 shows the time evolution of coordination numbers related to inter-molecular contacts between different types of molecular groups. The different colors indicate different values of the projected area per lipid (data for *A* = 1.4 *d*^2^ are shown in magenta; for *A* = 1.3 *d*^2^ in green; for *A* = 1.13 *d*^2^ in blue; and for *A* = 1.07 *d*^2^ in orange). The increase of *ζ*_*HB*_,*H*_*B*_ with time (figure 5A) implies that more inter-molecular contacts between head groups of lipid B is formed than broken during the simulation runs, reflecting clusterization of lipid B. The simultaneous decrease of *ζ*_*HB*_,*H*_*A*_ (figure 5B) is an effect of separation of lipid A from lipid B in the upper leaflet of the membrane. The increase of *ζ*_*HB*_,*P*_*C*_ with time (figure 5C) results from association of the polymer caps with the lipid-B head groups. The simultaneous increase of *ζ*_*PC*_,*P*_*C*_ (figure 5D) indicates that the polymers assemble at the membrane. The simultaneous formation of membrane-associated polymer clusters and segregation of lipids within the membrane resembles the adhesion-induced coalescence of nanoscale lipid clusters and adhesion receptors as studied recently in Monte Carlo simulations [29, 30].

In order to identify and characterize the assemblies of polymer P at the membrane surface, we performed clustering analysis of the configurations obtained from the DPD simulations (see Methods). Figure 6A shows the number *n*_*C*_ of clusters as a function of time. The different colors indicate different values of the projected area per lipid (data for *A* = 1.4 *d*^2^ are shown in magenta; for *A* = 1.3 *d*^2^ in green; for *A* = 1.13 *d*^2^ in blue; and for *A* = 1.07 *d*^2^ in orange). Here, we count only clusters that contain at least two polymers. In other words, single polymers that make no inter-molecular contacts are not considered as clusters.

**Figure 6:**
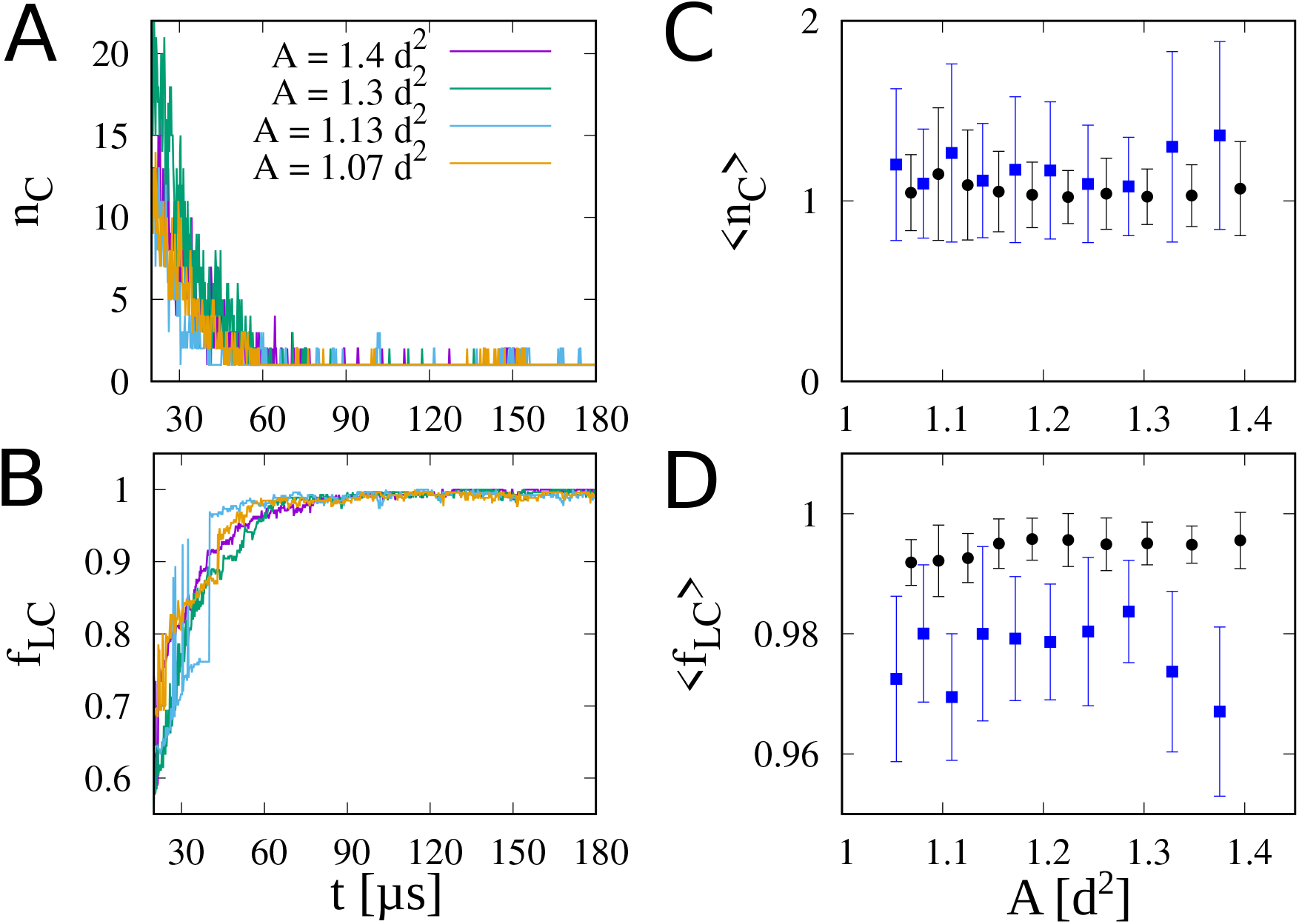
In the state of equilibrium, the polymers are coalesced into one cluster at the membrane surface. (A–B) The number *n*_*C*_ of clusters (A) and the fraction *f*_LC_ of polymers in the largest cluster (B) as a function of time. The different colors indicate different values of the projected area per lipid (data for *A* = 1.4 *d*^2^ are shown in magenta; for *A* = 1.3 *d*^2^ in green; for *A* = 1.13 *d*^2^ in blue; and for *A* = 1.07 *d*^2^ in orange). Here, the molar fraction of polymer P is *φ* = 10^−3^. (C–D) The average equilibrium values of *n*_*C*_ (C) and *f*_LC_ (D) as a function of the projected area per lipid *A*. The data points in blue and black correspond to the molar fractions *φ* = 5 *·* 10^−4^ and *φ* = 10^−3^, respectively. The error bars indicate standard deviation.

Figure 6B shows the fraction *f*_LC_ of polymers in the LC as a function time. The color code is as in igure 6A. Interestingly, in some trajectories *f*_LC_ is observed to change abruptly and discontinuously during equilibration (see, for example, the blue line in figure 6B). Such “jumps” in *f*_LC_ result from merging or breaking apart two clusters of comparable sizes, indicating that the polymer clusters are in a liquid state. However, after the equilibration of less than 90 *µ*s, there are only a couple of clusters in the system and practically all of the polymers are coalesced in the LC.

We averaged *n*_*C*_ and *f*_LC_ over the second halves of the DPD trajectories, i.e. over the time period from 90 to 180 *µ*s. The initial 90 *µ*s of equilibration was not used for the averaging. The resulting average values of *n*_*C*_ and *f*_LC_ at equilibrium are denoted as ⟨*n*_*C*_ ⟩ and ⟨*f*_LC_ ⟩and shown, respectively, in figures 6C and 6D as a function of *A*. The data points in blue and black correspond to the polymer molar fractions *φ* = 5 *·* 10^−4^ and *φ* = 10^−3^, respectively. The data shown in figures 6C and 6D together demonstrate that practically all of the polymers in the system at equilibrium form one large cluster, irrespective of the values of parameters *A* and *φ*. This polymer cluster at equilibrium will be referred to a the largest equilibrium cluster (LEC) hereafter.

### 3.4 Polymer nano-droplets form spontaneously on top of membrane regions curved inward

Within the clustering analysis we determined not only the number *N*_poly_ of the polymers forming the LEC, but also the number *N*_lip_ of the B-type lipids in contact with the LEC. The ratio *N*_lip_*/N*_poly_ averaged over the second halves of the DPD trajectories (i.e. over the time window from 90 to 180 *µ*s) is denoted as ⟨*N*_lip_*/N*_poly_ ⟩ and shown in figure 7A as a function of *A*. The data points in blue and black correspond to the molar fractions *φ* = 5 *·* 10^−4^ and *φ* = 10^−3^, respectively. Interestingly, ⟨*N*_lip_*/N*_poly_ ⟩ decreases monotonically with *A*, i.e. takes smaller values for planar membranes than for bent membranes. This means that more lipids of type B are recruited to the polymer LEC when the membrane is bent, indicating that membrane curvature can be a relevant factor in effective interactions between the polymer nano-droplet and the lipid membrane.

**Figure 7:**
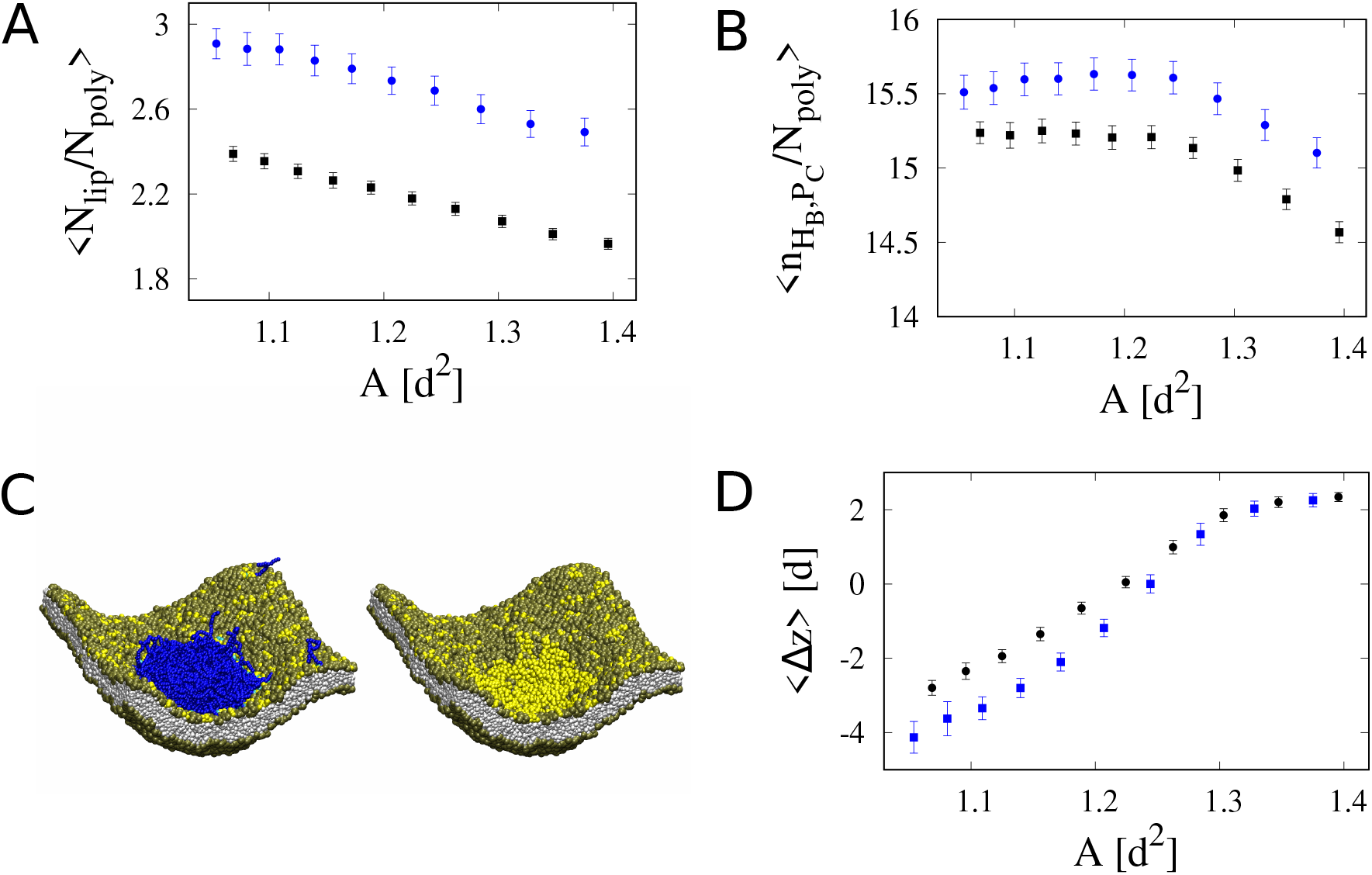
The polymer nano-droplets localize on top of membrane regions that are curved inward. (A) ⟨*N*_lip_*/N*_poly_ ⟩as a function of *A* for *φ* = 5 *·* 10^−4^ (points in blue) and *φ* = 10^−3^ (points in black). *N*_poly_ is the number of the polymers forming the LEC, *N*_lip_ is the number of the B-type lipids directly contacting the LEC, and the angular brackets ⟨… ⟩ represent the equilibrium average. (B) ⟨*n*_*HB*_,*P*_*C*_ */N*_poly_ ⟩ as a function of *A*, where *n*_*HB*_,*P*_*C*_ denotes the number of contacts between the B-type head-group beads *H*_*B*_ and the cap beads *P*_*C*_ of the polymers forming the LEC (color code as in panel A). (C) Two simulation snapshots taken at *t*=180 *µ*s. Here, *A* = 1.1 *d*^2^ and *φ* = 10^−3^. The water beads *W* are not shown for clarity. The snapshot on the left hand side shows that polymers P form a nano-droplet inside a ‘valley’ or a ‘pit’ on the membrane surface. In the snapshot on the right hand side, only the lipids are shown. It can be seen that the B-type lipids (yellow) are segregated from the A-type lipids (tan) and localized below the polymer nano-droplet and in the membrane region curved inward. (D) The equilibrium average of the nano-droplet immersion depth, ∆*z*, as a function of *A* (color code as in panels A and B).

We also analyzed the number *n*_*HB*_,*P*_*C*_ of contacts between the B-type head-group beads *H*_*B*_ and the cap beads *P*_*C*_ of the polymers forming the LEC. Figure 7B shows the average number of these *H*_*B*_-*P*_*C*_ contacts per polymer, ⟨*n*_*HB*_,*P*_*C*_ */N*_poly_ ⟩, as a function of area per lipid. Note that the number of *H*_*B*_-*P*_*C*_ contacts between a lipid B and a polymer P can not exceed 16, because one lipid of type B contains four head-group beads *H*_*B*_ and one polymer comprises four cap beads *P*_*C*_. The data shown in figures 7A and 7B together mean that the total number of contacts between the polymer LEC and the B-type lipid heads decreases with increasing the area per lipid, implying that the polymer droplet adheres tighter to bent membranes with *A <* 1.2 *d*^2^ than to flat membranes with *A >* 1.2 *d*^2^.

Visual inspection of the simulation system configurations reveals that the LECs are localized in ‘valleys’ or ‘pits’ on the membrane surface, as depicted in the simulation snapshot in figure 7C. To quantify this observation we introduce an ‘immersion depth’ of the nano-droplet, defined as ∆*z* = *z*_2_ −*z*_1_, where *z*_2_ is the z-coordinate of the center of mass of all the lipid-tail beads (i.e. *T* -type beads) and *z*_1_ is the z-coordinate of the center of mass of the lipid-binding beads (i.e. *P*_*C*_-type beads) of the polymers forming the LEC. Figure 7D shows ⟨∆*z* ⟩as a function of *A*, where the angular brackets denote the average over the second halves of the DPD trajectories (i.e. over the time window from 90 to 180 *µ*s), approximating thus the equilibrium average. Interestingly, ⟨∆*z* ⟩is found to decrease monotonically with *A*. In the case of planar membranes with larger area-per-lipid values, ⟨∆*z* ⟩*>* 0. In the case of bent membranes with smaller area-per-lipid values, ⟨∆*z* ⟩ *<* 0. Taken together, the data shown in figure 7D mean that the more the membrane is bent, the lower in the valley the nanodroplet is located. This result implies that the nano-droplets of polymer P sense the concavity of the membrane surface, despite the fact the single polymers have no particular shape or specific curvature that could possibly match the membrane curvature.

### 3.5 A toy-model can rationalize the sensing of membrane curvature by liquid droplets

To better understand the sensing of membrane curvature by liquid droplets, we examined a toy model illustrated in figure 8A: A droplet is placed on a membrane segment shaped like a spherical cap with radius *R*. The contact angle *θ* between the liquid-vapor and liquid-membrane interfaces at equilibrium is given by the Young’s equation, cos *θ* = (*σ*_vm_ − *σ*_lm_) */σ*, where *σ*_vm_ and *σ*_lm_ are tensions of the vapor-membrane and liquid-membrane interfaces, respectively, and *σ* is the tension of the liquid-vapor interface. Thus the contact angle *θ* is determined solely by the three surface tensions, and not by the geometry of the model.

**Figure 8:**
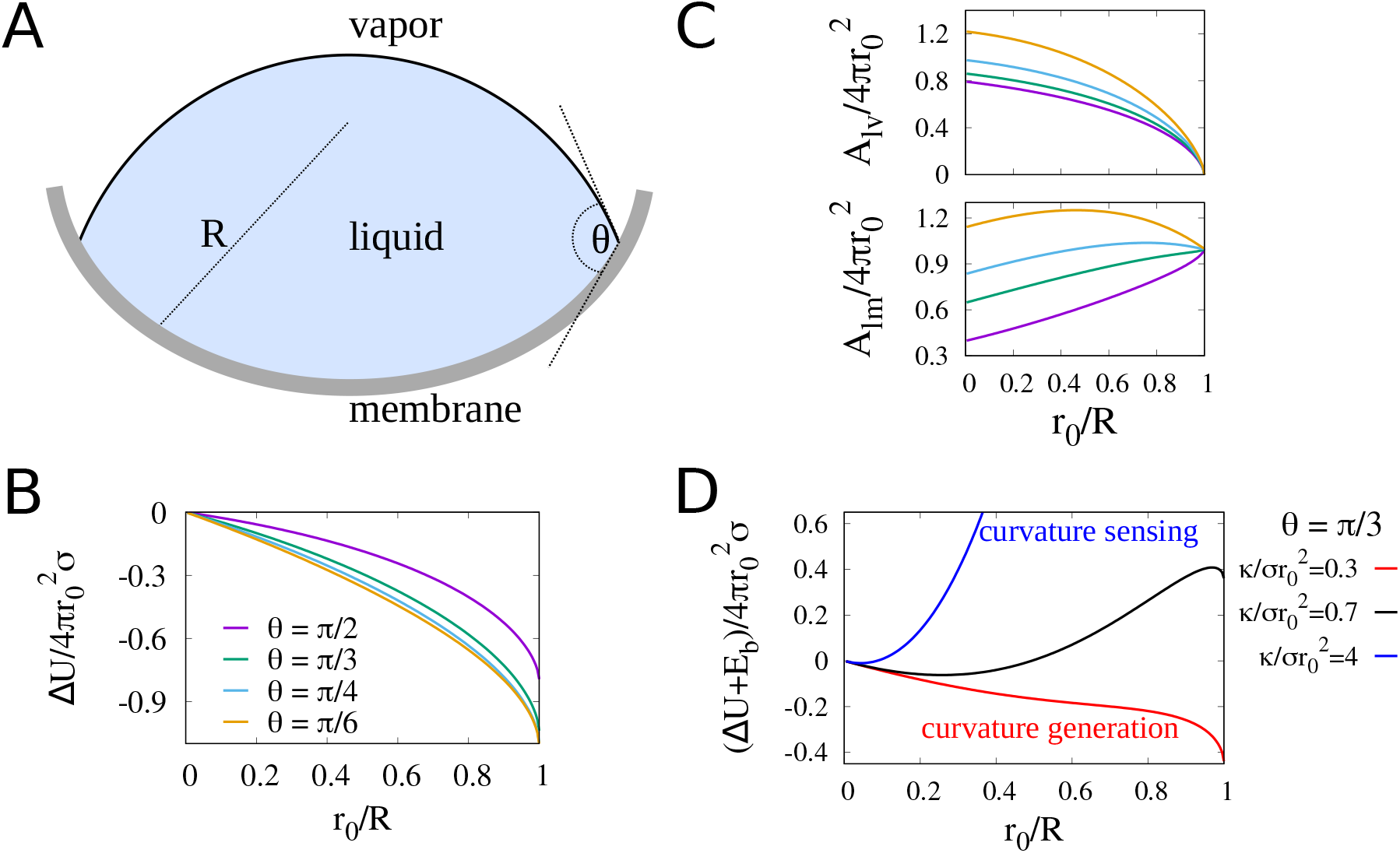
(A) Illustration of the spherical cap model. The membrane has the shape of a spherical cap with radius *R*. The contact angle *θ* is given by the Young’s equation and, thus, determined solely by surface tensions of the liquid-vapor, liquid-membrane and vapor-membrane interfaces. (B) The difference in the interfacial energy between curved and planar membrane configurations, ∆*U* = *U* (*r*_0_*/R, θ*) *U* (0, *θ*), as a function of the dimensionless curvature *r*_0_*/R* of the membrane. ∆*U* is given in units of 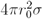, where *σ* denotes the liquid surface tension and *r*_0_ defines the droplet volume 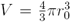. The curves in different colors represent results obtained for different values of the contact angle (data for *θ* = *π/*6 in orange; for *θ* = *π/*4 in blue; for *θ* = *π/*3 in green; and for *θ* = *π/*2 in purple). (C) The areas of the liquid-vapor and liquid-membrane interfaces, *A*_lv_ and *A*_lm_, as a function of the dimensionless curvature *r*_0_*/R* of the membrane. The areas are given in units of the area 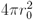 a spherical droplet of volume 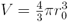. The color code is as in panel B. (D) The total energy ∆*U* +*E*_b_ of the droplet and the membrane as a function of *r*_0_*/R* for *θ* = *π/*3 and 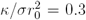 (red), 0.7 (black) and 4 (blue). *E*_b_ is the energy of bending the membrane from the planar configuration to the spherical cap with radius *R*, and *κ* denotes the bending rigidity modulus of the membrane.

Next, we assume that the droplet has a fixed volume *V*. A spherical droplet of the same volume has radius *r*_0_ given by 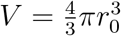 We use the dimensionless parameter *r*_0_*/R* as a measure of curvature of the membrane.

The interfacial energy *U* of the system shown in figure 8A can be calculated exactly and expressed as a function of *r*_0_*/R* and *θ* (see Supplementary Material). Figure 8B shows the difference in the interfacial energy between curved and planar membrane configurations, ∆*U* = *U* (*r*_0_*/R, θ*) − (0, *θ*), as a function of *r*_0_*/R* for different values of *θ*. Here, ∆*U* is given in units of 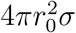 i.e. the surface energy of a free spherical droplet with radius *r*_0_.

The calculation results shown in figure 8B mean that the interfacial energy of the droplet-membrane system decreases as the membrane gets more concave, independent of the contact angle. This result is consistent with the capillary effect of curvotaxis [32, 14, 34] and appears to be in agreement with the sensing of membrane curvature by nano-droplets as observed in the DPD simulations.

Figure 8C shows the areas of the liquid-vapor and liquid-membrane interfaces, *A*_lv_ and *A*_lm_, as functions of *r*_0_*/R* for different values of *θ*. The color code is as in figure 8B. The areas are given in units of 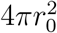 i.e. the surface area of a free spherical droplet with radius *r*_0_. Interestingly, *A*_lv_ decreases with *r*_0_*/R* whereas *A*_lm_ increases with *r*_0_*/R*. Thus, the decrease in the interfacial energy ∆*U* with increasing the membrane curvature results from shrinking the liquid-vapor interface and expanding the liquid-membrane contact.

If the decrease in the interfacial energy ∆*U* is larger than the energy *E*_b_ of bending the membrane from the planar configuration to the spherical cap with radius *R*, then the interfacial tensions shall cause the membrane to bend. Otherwise, the droplet senses the membrane curvature (∆*U <* 0) but can not induce membrane bending (∆*U* + *E*_b_ *>* 0). Figure 8D shows the total energy of the droplet-membrane system, ∆*U* + *E*_b_, as a function of *r*_0_*/R* for *θ* = *π/*3 and different values of a dimensionless parameter 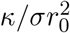 where *κ* denotes the bending rigidity modulus of the membrane. For 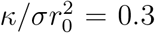 (curve in red), ∆*U* + *E*_b_ is negative and monotonically decreases with *r*_0_*/R*, corresponding to the case of curvature generation. For 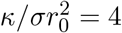 (curve in blue), ∆*U* + *E*_b_ increases with *r*_0_*/R*, corresponding to the case of curvature sensing. For 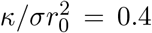 (curve in black), ∆*U* + *E*_b_ is a non-monotonic function of *r*_0_*/R* with the minimum at *r*_0_*/R ≈* 0.25, which means that the membrane can be bent by the droplet only to some extent. The analysis illustrated in figure 8D implies that larger droplets with sufficient surface tension *σ* are capable of bending flexible membranes, or generating membrane curvature, whereas smaller droplets can function as sensors of membrane curvature.

## 4 Discussion

Membrane curvature sensing by folded proteins with certain structural motifs has been studied in details [1]. Crescent-shaped BAR (Bin/Amphiphysin/Rvs) domain proteins bind selectively to highly curved membranes by their concave side [40]. Amphipathic helices and curved scaffolds are common motifs found in proteins participating in clathrin-mediated endocytosis [27]. However, intrinsically disordered regions (IDRs) are also highly abundant within the same proteins. The possibility that IDRs contribute to membrane curvature sensing seems to be largely overlooked, probably because of the common concept that curvature sensing requires specific structural motifs.

Increasingly more evidence suggests that in spite of lacking any specific structural motifs, IDRs and IDPs can be potent sensors of membrane curvature [60]. Two mechanisms have been identified to account for the membrane curvature sensing by IDPs [59]. The first one is entropic in nature: IDPs bind preferentially to membrane regions curved outward because the more convex the membrane is, the larger the conformational entropy of membrane-associated IDPs becomes. The second mechanism arises in cases when both the membrane surface and the IDP are negatively charged: When the IDP is bound to the membrane, the electrostatic repulsion between anionic amino acid residues and lipid head groups is weaker for membranes curved outward than for flat membranes, simply because the average distance between the two groups of objects gets larger when the membrane curvature increases.

In contrast to the aforementioned mechanisms, which explain why IDPs bind preferentially to convex membranes, our simulations demonstrate that IDP nano-droplets get localized in membrane pits or valleys, where the convex shape of the nano-droplet fits the concave surface of the membrane. If the IDP lipid-binding sites are exposed mostly on the surface of the nano-droplet, as in our simulations (figure 7), the nano-droplet can make more energetically favorable contacts with a concave membrane surface than with a flat membrane, which is the driving force for membrane curvature sensing in our model.

Our results of the coarse-grained simulations (section 3.4) and analytical calculations (section 3.5) indicate together towards a generic mechanism of membrane curvature sensing, which we expect to be independent of molecular level details of IDPs forming nano-droplets or BCs. As a matter of fact, such molecular features of IDPs as their amino acid sequence and local secondary structure elements are not taken into account in the coarse-grained polymer model studied here. Still, an important extension of our work will be to study membrane curvature sensing by BCs using coarse-grained models that take into account the amino acid sequence of IDPs [13, 37, 52].

Membrane bending by protein phase separation has been recently observed in microscopy experiments [57]. The corresponding mechanism of membrane curvature generation has been attributed a net compressive stress on one side of the membrane as created by the protein phase separation at the membrane surface. In contrast, in the coarse-grained simulations reported here, the nano-droplets were not observed to induce membrane bending. Instead, when the membrane was kept compressed, it was bent independent of interactions with the IDPs, and then the nano-droplets were found to form spontaneously on top of membrane regions curved inward. The toy model presented in section 3.5 illustrates that the switch between curvature generation and curvature sensing is determined by an interplay of the membrane bending rigidity and the droplet interfacial energy.

An important objective for future studies would be to employ the DPD coarse-grained model to explore mechanisms of membrane curvature generation by IDPs and their BCs. Such future studies can in particular help to identify the mechanism of endocytic pit formation by BCs of galectin-3. On the clathrin mediated endocytic pathway, proteins with specific shapes (such as epsins and BAR-domain proteins) impose their intrinsic curvature on the membrane. Since galectin-3 is an IDP, it likely employs a different mechanism to deform membranes. We anticipate that the endocytic pit formation mediated by galectin-3 is a collective and cooperative process, in which the specific binding of galectin-3 to glycosphingolipids and the condensation of galectin-3 occur simultaneously, resulting in stabilization or generation of membrane pits.

Recent studies have shown that mechanical properties of lipid membranes are strongly affected by lipid composition asymmetry (curvature stress) and lipid number asymmetry (differential stress) [35, 21]. Therefore, another important direction of future studies employing the DPD coarse-grained model would be to investigate how these asymmetries influence membrane curvature generation processes.

## Supporting information

Supplementary Material: Exact solution of the toy model presented in section 3.5

## Author Contributions

Midhun Mohan Anila: data curation, formal analysis, investigation, writing – original draft. Rikhia Ghosh: formal analysis, writing – original draft, writing – review and editing. Bartosz Ró ż ycki: conceptualization, data curation, formal analysis, funding acquisition, investigation, methodology, resources, supervision, visualization, writing – original draft, writing – review and editing.

## Acknowledgements

The work has been financially supported by the National Science Centre of Poland via grant number 2020/39/B/NZ1/00377. The simulations were carried out using the supercomputer resources at the Centre of Informatics – Tricity Academic Supercomputer and networK (CI TASK) in Gda ń sk, Poland.

## Footnotes

1 available at https://github.com/Osprey-DPD/osprey-dpd

