## Supplementary Material: Exact solution of the toy model presented in section 3.5 for "Membrane curvature sensing by model biomolecular condensates"

### Supplementary Material
#### Membrane curvature sensing by model biomolecular condensates

Midhun Mohan Anila[a], Rikhia Ghosh[b], and Bartosz Różycki[a*]
[a]Institute of Physics, Polish Academy of Sciences,
Al. Lotników 32/46, 02-668 Warsaw, Poland
[b]Icahn School of Medicine at Mount Sinai,
1 Gustave L. Levy Pl, New York, NY 10029, USA

We consider a spherical cap model illustrated in figure 1. A liquid droplet is placed inside a sphere of radius $R$. Alternatively one can think of a droplet placed on a concave membrane that is shaped as a spherical cap with radius $R$. The liquid-vapor interface is modeled as a spherical cap with radius $r$. The contact angle $\theta$ between the liquid-vapor and liquid-membrane interfaces is given by the Young's equation

$$\cos\theta = \frac{\sigma_{\rm vm} - \sigma_{\rm lm}}{\sigma} \tag{1}$$

where $\sigma$, $\sigma_{\rm lm}$ and $\sigma_{\rm vm}$ denote the surface tensions of the liquid-vapor, liquid-membrane and vapor-membrane interfaces. On the other hand, the geometry of the system illustrated in figure 1 implies that

$$\theta = \alpha + \beta \tag{2}$$

and

$$R \sin\alpha = r \sin\beta \tag{3}$$

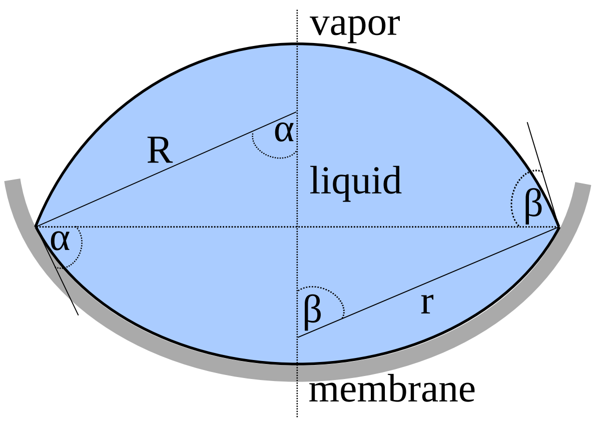


Figure 1: Illustration of the spherical cap model. The liquid-vapor interface is a spherical cap with radius $r$. The membrane has the shape of a spherical cap with radius $R$. The contact angle $\theta = \alpha + \beta$ is given by the Young's equation (1), and $R\sin\alpha = r\sin\beta$.

*

The volume of the droplet can be written as

$$V = \frac{4}{3}\pi R^3 g(\alpha) + \frac{4}{3}\pi r^3 g(\beta) \tag{4}$$

where

$$g(\alpha) = \frac{1}{4}\left(2 + \cos\alpha\right)\left(1 - \cos\alpha\right)^2 \tag{5}$$

A spherical droplet with the same volume $V$ has radius $r_0$ given by

$$V = \frac{4}{3}\pi r_0^3 \tag{6}$$

Therefore, using equations (2) and (3), we obtain

$$\left(\frac{r_0}{R}\right)^3 = g(\alpha) + g(\theta - \alpha)\frac{\sin^3(\alpha)}{\sin^3(\theta - \alpha)} \tag{7}$$

Equation (7) determines the angle $\alpha$ for given contact angle $\theta$ and dimensionless curvature $r_0/R$ of the surface.

The total interfacial energy of the system illustrated in figure 1 is

$$U = A_{\mathrm{lv}}\sigma + A_{\mathrm{lm}}\left(\sigma_{\mathrm{lm}} - \sigma_{\mathrm{vm}}\right) = \sigma\left(A_{\mathrm{lv}} - A_{\mathrm{lm}}\cos\theta\right) \tag{8}$$

where $A_{\mathrm{lm}}$ and $A_{\mathrm{lv}}$ denote the areas of the liquid-membrane and liquid-vapor interfaces, respectively, and the second equality in (8) follows from the Young's equation. Within the framework of the spherical cap model, these areas are given by

$$A_{\mathrm{lm}} = 2\pi R^2\left(1 - \cos\alpha\right) \tag{9}$$

and

$$A_{\mathrm{lv}} = 2\pi r^2\left(1 - \cos\beta\right) = 2\pi R^2\left(1 - \cos(\theta - \alpha)\right)\frac{\sin^2(\alpha)}{\sin^2(\theta - \alpha)} \tag{10}$$

where the second equality in (10) follows from equations (2) and (3). It is convenient to use dimensionless areas of the two interfaces

$$\frac{A_{\mathrm{lm}}}{4\pi r_0^2} = \frac{1}{2}\left(\frac{R}{r_0}\right)^2\left(1 - \cos\alpha\right) \tag{11}$$

and

$$\frac{A_{\mathrm{lv}}}{4\pi r_0^2} = \frac{1}{2}\left(\frac{R}{r_0}\right)^2\left(1 - \cos(\theta - \alpha)\right)\frac{\sin^2(\alpha)}{\sin^2(\theta - \alpha)} \tag{12}$$

where $4\pi r_0^2$ is the area of a spherical droplet with volume $V = \frac{4}{3}\pi r_0^3$. Now the total interfacial energy of the droplet, as given by equation (8), can be now written as

$$\frac{U}{4\pi r_0^2\sigma} = \frac{1}{2}\left(\frac{R}{r_0}\right)^2\left(1 - \cos(\theta - \alpha)\right)\frac{\sin^2(\alpha)}{\sin^2(\theta - \alpha)} - \frac{1}{2}\left(\frac{R}{r_0}\right)^2\left(1 - \cos\alpha\right)\cos\theta \tag{13}$$

where $4\pi r_0^2\sigma$ is the interfacial energy of a free, spherical droplet of radius $r_0$. In the limiting case of a flat substrate, i.e. $R \to \infty$ and $\alpha \to 0$ with $R\alpha = r\sin\theta$, we obtain

$$\frac{U}{4\pi r_0^2\sigma} = \left(g(\theta)\right)^{1/3} \tag{14}$$

Thus the difference in the interfacial energy of the droplet placed on the curved and planar membrane, $\Delta U = U(\theta, r_0/R) - U(\theta, 0)$, is given by

$$\frac{\Delta U}{4\pi r_0^2 \sigma} = \frac{1}{2}\left(\frac{R}{r_0}\right)^2 (1-\cos(\theta-\alpha))\frac{\sin^2(\alpha)}{\sin^2(\theta-\alpha)} - \frac{1}{2}\left(\frac{R}{r_0}\right)^2 (1-\cos\alpha)\cos\theta - (g(\theta))^{1/3} \quad (15)$$

By solving equation (7) for $\alpha$ and substituting to equation (15) we obtain $\Delta U$ as a function of the Young's contact angle $\theta$ and the dimensionless curvature $r_0/R$ of the membrane. Figure 8B in the main text shows $\Delta U$ in units of $4\pi r_0^2\sigma$ as a function of $r_0/R$ for four different values of the contact angle $\theta$. Interestingly, independent of the $\theta$-value, $\Delta U$ is found to decrease with increasing curvature of the membrane. The droplet thus 'feels' the curvature of the membrane.

To clarify why $\Delta U$ is a decreasing function of $r_0/R$, we use equations (7), (11) and (12) to determine the areas of the liquid-vapor and liquid-membrane interfaces as functions of $\theta$ and $r_0/R$. Figure 8C in the main text shows that $A_{\mathrm{lv}}$ decreases and $A_{\mathrm{lm}}$ increases with $r_0/R$. This means that the decrease in $\Delta U$ with increasing $r_0/R$ is caused by shrinking the liquid-vapor interface and expanding the liquid-surface interface. Intuitively it seems clear that the energetically favorable contact between the droplet and the membrane gets larger as the membrane gets more concave.

The total energy of the system is a sum of the droplet interfactal energy $\Delta U$ and the energy of bending the membrane

$$E_{\mathrm{b}} = 2\kappa \int M^2 \mathrm{d}A = \frac{2\kappa}{R^2} A_{\mathrm{lm}} \quad (16)$$

Here $\kappa$ is the bending rigidity modulus of the membrane and $M = 1/R$ is the mean curvature. Equations (11) and (16) lead to

$$\frac{E_{\mathrm{b}}}{4\pi r_0^2 \sigma} = \frac{\kappa}{\sigma r_0^2}(1-\cos\alpha) \quad (17)$$

By solving equation (7) for $\alpha$ and substituting to equation (17) we obtain the energy $E_{\mathrm{b}}$ of bending the membrane from a flat state to a spherical cap with radius $R$.

Figure 8D in the main text shows the total energy $\Delta U + E_{\mathrm{b}}$ in units of $4\pi r_0^2\sigma$ as a function of $r_0/R$ for $\theta = \pi/3$ and three values of the dimensionless parameter $\kappa/\sigma r_0^2$. Interestingly, for small values of $\kappa/\sigma r_0^2$ (see the red curve in figure 8D), the total energy $\Delta U + E_{\mathrm{b}}$ decreases with the membrane curvature $r_0/R$, implying that the droplet-membrane intercations drive, or generate, membrane curvature. For large values of $\kappa/\sigma r_0^2$ (see the blue curve in figure 8D), on the other hand, $\Delta U + E_{\mathrm{b}}$ increases while $\Delta U$ decreases with increasing $r_0/R$, implying curvature sensing rather than curvature generation. For intermediate values of $\kappa/\sigma r_0^2$ (see the black curve in figure 8D), $\Delta U + E_{\mathrm{b}}$ has a minimum at a certain curvature $r_0/R$.
